# A cell-type-specific multi-protein complex regulates expression of Cyclin B protein in *Drosophila* male meiotic prophase

**DOI:** 10.1101/2023.02.16.528869

**Authors:** Catherine C. Baker, Lorenzo Gallicchio, Lucineh Parsanian, Emily Taing, Cheuk Tam, Margaret T. Fuller

**Affiliations:** Department of Developmental Biology, Stanford University School of Medicine, Stanford, CA, 94305, USA

**Keywords:** *Drosophila*, spermatogenesis, meiosis, translation, RNA, Cyclin B

## Abstract

During meiosis, germ cell and stage-specific components impose additional layers of regulation on the core cell cycle machinery to set up an extended G2 period termed meiotic prophase. In *Drosophila* males, meiotic prophase lasts 3.5 days, during which spermatocytes turn up expression of over 3000 genes and grow 25-fold in volume. Previous work showed that the core cell cycle regulator Cyclin B (CycB) is subject to translational repression in immature *Drosophila* spermatocytes, mediated by the RNA-binding protein Rbp4 and its partner Fest. Here we show that another spermatocyte-specific protein, Lut, is required for translational repression of *cycB* in an 8-hour window just before spermatocytes are fully mature. In males mutant for *rbp4* or *lut*, spermatocytes enter and exit the meiotic divisions 6-8 hours earlier than in wild-type. In addition, we show that spermatocyte-specific isoforms of Syncrip (Syp) are required for expression of CycB protein and normal entry into the meiotic divisions. Both Lut and Syp interact with Fest in an RNA-independent manner. Thus a complex of spermatocyte-specific regulators choreograph the timing of expression of CycB protein during male meiotic prophase.

**SUMMARY STATEMENT:** Expression of a conserved cell cycle component, Cyclin B, is regulated by multiple mechanisms in the *Drosophila* male germline to dictate the correct timing of meiotic division.

## INTRODUCTION

A key aspect of specialization within a given tissue or stem cell lineage is regulation of the cell cycle: when to divide and if so, whether to divide symmetrically or asymmetrically, whether to exit the cell cycle completely or to undertake unusual cell cycles. Unconventional cell cycles include those that skip M phase leading to polyploidy, and meiosis, which features an extended G2 cell cycle phase termed meiotic prophase. Both extrinsic cues and intrinsic, lineage-specific factors feed into regulation of the core cell cycle machinery to generate cell-type and tissue-specific control of the cell cycle. For example, the mechanisms that regulate the length of meiotic prophase in females have been extensively studied and are strongly influenced by extrinsic factors such as hormonal control. In human females, meiotic prophase can last 12-50 years. However, less is known about the mechanisms that regulate meiotic prophase in males, where it tends to last a specified time depending on the species: 12 days in mouse and 3.5 days in *Drosophila*.

Studies of the *Drosophila* male germline are beginning to reveal how intrinsic factors controlled by the male germ cell developmental program impose layers of regulation on core cell cycle machinery components during the specialized cell cycle of meiosis in males. In the *Drosophila* testis, germ cells are arrayed in a roughly spatiotemporal order originating from germline stem cells at the apical tip of the testis (Fig. 1A). When germline stem cells divide they leave a stem cell daughter at the hub and release a daughter cell (gonialblast) that will give rise to all of the later germ cell types. The gonialblast initiates four rounds of synchronous mitotic divisions with incomplete cytokinesis, producing 16 interconnected germ cells that undergo pre-meiotic S phase. During the subsequent meiotic prophase, the resulting spermatocytes grow 25-fold in volume and turn on the massive transcription program required for post-meiotic spermatid differentiation. The 3.5-day meiotic G2 phase is characterized by developmental mechanisms that delay the appearance of core cell cycle regulatory proteins such as Cyclin B (CycB) and the Cdc25 phosphatase Twine (Twe) until just before they are needed for the meiotic divisions (White-Cooper et al., 1998; Maines and Wasserman, 1999).

**Figure 1.**
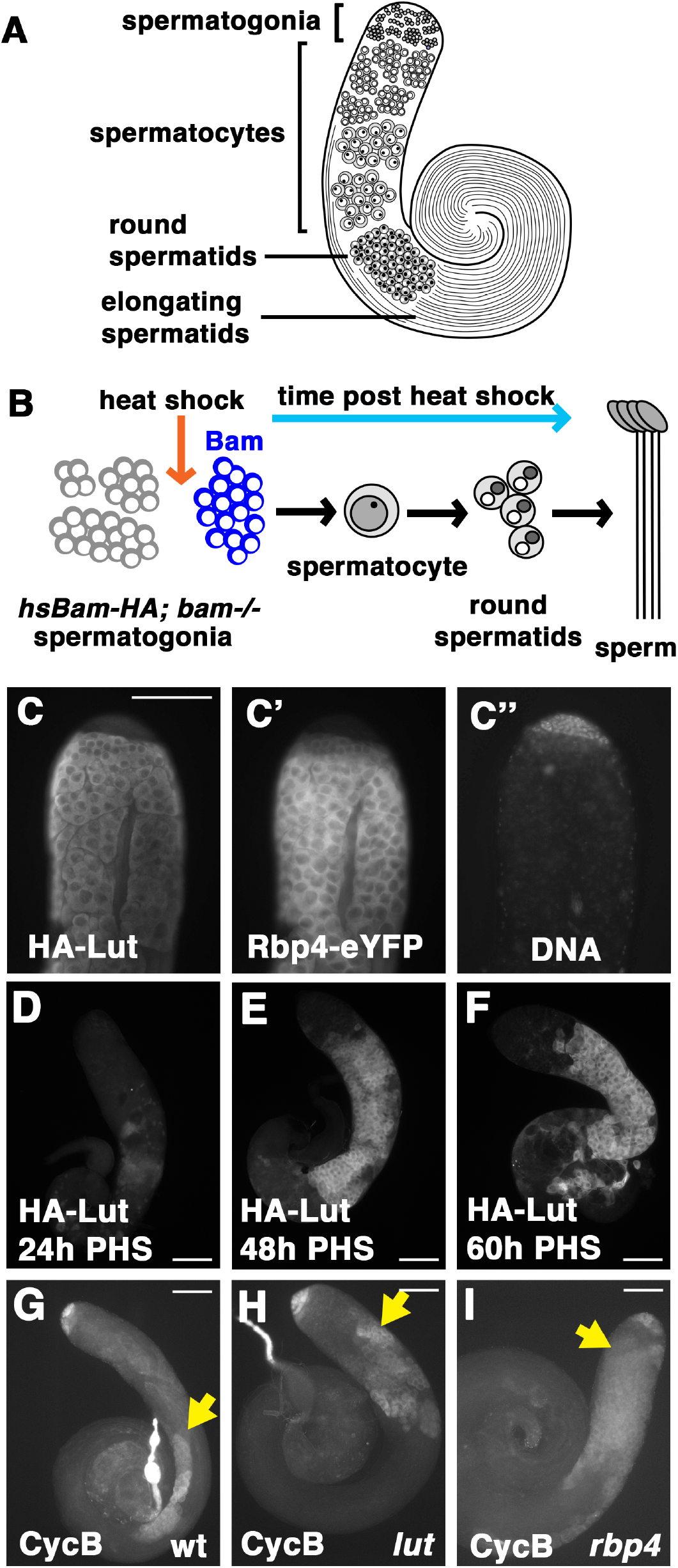
Lutin is expressed early in spermatocyte development and is required for repression of *cycB* translation in mid-stage spermatocytes. (A) Illustration of a wild-type testis with major germline cell types labeled. (B) Schematic of the hs-Bam differentiation time-course. (C) Immunofluorescence image of the apical tip of a testis from a male carrying HA-Lut and Rbp4-eYFP transgenes, stained with (C) anti-HA (C’) anti-GFP (C’’) DAPI. (D-F) Anti-HA staining of apical tips of *bam−/− hsBam HA-Lut* testes collected at (D) 24h post-heat-shock (PHS), (E) 48h PHS (F) 60h PHS. (G-I) Anti-CycB staining of (G) wt (H) *lut* and (I) *rbp4* testes. Yellow arrows indicate onset of CycB protein expression in spermatocytes. Scale bar: 100 μm for C-I.

Here we show that a complex of cell-type-specific proteins and protein isoforms bound to the 3’UTR of the *cycB* RNA regulate the timing of CycB protein expression during male meiotic prophase in *Drosophila*. We had previously shown that the RNA-binding protein Rbp4 and its binding partner Fest were required to repress translation of *cycB* in immature spermatocytes. Rbp4 bound the 130 nt *cycB* 3’UTR expressed in spermatocytes in biotin pulldown assays, suggesting that Rbp4 might act directly on the *cycB* RNA to block its translation (Baker et al., 2015). We now show that the novel protein Lutin (encoded by CG1690), is co-expressed at the onset of spermatocyte differentiation with Rbp4 and Fest and interacts with Fest (and with Rbp4, via Fest) independently of RNA. Lut is also required for cell-type-specific translational repression of *cycB* in spermatocytes, although at a later stage. CycB protein appears about 50 hours early in spermatocytes of males mutant for *rbp4*, but only about 8 hours early in males mutant for *lut*. Spermatocytes mutant for either *rbp4* or *lut* enter and exit the meiotic divisions about 6-8 hours earlier than in wild type, indicating that expression of CycB protein can be permissive for meiotic entry in late-stage spermatocytes. We also show that testis-specific isoforms of Syp, the *Drosophila* homolog of mammalian hnRNP Q, are required for expression of CycB protein in mature spermatocytes, as well as for normal meiotic progression and post-meiotic differentiation. A cytoplasmic isoform of Syp expressed in spermatocytes binds the 130 nt *cycB* 3’UTR and interacts with the repressive Rbp4-Fest-Lut complex via Fest throughout spermatocyte development. Data from the *syp* mutant suggest a role for Syp in stabilizing the *cycB* RNA, and results from *rbp4 syp* and *lut syp* double mutants point to some possible models for how the three proteins might act in series or in parallel to regulate *cycB* RNA stability and translation during the extended meiotic G2 prophase.

## RESULTS

### Lutin represses premature translation of *cycB* in spermatocytes

The *Drosophila* gene CG1690 encodes a 15kD predicted protein that has no currently recognized domains, but is conserved in most species within the *Drosophila* genus, notably the entire subgenus *Sophophora* as well as Hawaiian *Drosophila*. The only homolog found outside this genus is in *Scaptodrosophila lebanonensis*. We named CG1690 *lutin* after the small and pleasantly-mischievous French house-elf. Expression of CG1690/Lutin initiates early in spermatocyte development, as first revealed by RNA-seq data from a heat-shock Bam (hs-Bam) differentiation time-course (Lu et al., 2020). This technique exploits the fact that *bam* mutant testes accumulate spermatogonia, but following a short pulse of wild-type Bam expression under the control of a heat-shock promoter, germ cells march through differentiation in rough synchrony (Kim et al., 2017 and illustrated in Fig. 1B). RNA-seq analysis of testes from *hsBam; bam* flies subjected to a single pulse of heat shock showed that *lutin* RNA was upregulated over 1000-fold by 48 hours post-heat-shock (PHS) compared to testes from *bam* flies heat-shocked and cultured in parallel. A transgene composed of the endogenous *lut* promoter driving a *lut* cDNA with a 3xHA N-terminal tag showed expression of HA-Lut protein starting in very early spermatocytes (Fig. 1C), in a pattern very similar to Rbp4-eYFP (Fig. 1C’). Like Rbp4-eYFP, the HA-Lut protein was cytoplasmic. Consistent with the time-course RNA-seq data, expression of HA-Lut from the transgene was not detected in germ cells at 24h PHS but was robustly expressed in differentiating spermatocytes at 48h and 60h PHS (Fig. 1D-F).

Function of Lutin is required to repress premature expression of CycB protein in spermatocytes. A null allele of *lut* made by CRISPR (Materials and Methods) carried a 5 nt microdeletion of nucleotides 136-140 of the *lut* protein coding sequence (*lut^1^*). This resulted in a frameshift just after the codon for Gly45, followed by three novel residues (I, W, G), then a stop codon, leading to a Lut protein missing nearly two thirds of its sequence.

Immunofluorescence staining revealed expression of CycB protein in spermatocytes closer to the apical tip of the testis in *lut^1^/Df* males (hereafter referred to as *lut*) than in wild type (Fig. 1H vs. 1G), although not as apical (early) as seen in testes from males mutant for *rbp4* (Fig. 1I).

### Lut interacts with Rbp4 via binding to Fest

The Lut protein physically interacts with Fest, independently of RNA. Immunoprecipitation of eYFP-Fest with anti-GFP from testis extracts brought down HA-Lut, both in the presence or absence of RNAse A (Fig. 2A, right side). In the reciprocal experiment, immunoprecipitation of V5-Lut from testis extracts in the presence of RNAse brought down HA-Fest (Fig. 2B). Immunoprecipitation of HA-Lut with eYFP-Fest held true even in testes mutant for the Fest-interacting protein Rbp4. In contrast, although immunoprecipitation of Rbp4-eYFP brought down HA-Lut from wild-type extracts in the presence and absence of RNAse (Fig. 2A, left side), HA-Lut did not co-immunoprecipitate with Rbp4 in a *fest* mutant background, indicating that Rbp4 requires Fest for its interaction with Lut. Together these data suggest a model where Rbp4, Fest, and Lut form a repressive complex in which Fest acts as a scaffold. As Baker et al. (2015) showed that Rbp4 protein binds the *cycB* 3’UTR, Rbp4 may thus recruit the repressive complex to the *cycB* RNA (Fig. 2C).

**Figure 2.**
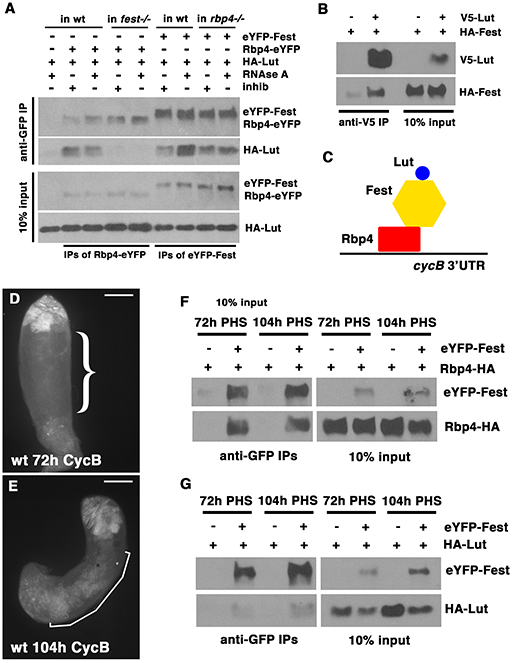
Lutin interacts with Fest, and with Rbp4 dependent on Fest. (A) Western blots probed with anti-GFP or anti-HA showing proteins immunoprecipitated with anti-GFP from testis extracts from flies expressing HA-Lut and either Rbp4-eYFP (lanes 2-5) or eYFP-Fest (lanes 6-9). Negative control: HA-Lut alone (lane 1). RNAse A or RNAse inhibitor was added as indicated. Flies used in lanes 4-5 were *fest* mutants. Flies used in lanes 8-9 were *rbp4* mutants. Western blots probed with anti-V5 or anti-HA of anti-V5 immunoprecipitates of testis extracts from flies expressing V5-Lut and HA-Fest. RNase A added to all samples. Negative control: HA-Fest alone. (C) Schematic of proposed repressive complex, with Lut and Rbp4 both binding to Fest. (D,E) Testes from the hs-Bam time-course immunostained with anti-CycB. Scale bars: 100 μm. (D) 72h PHS. Curly bracket: region of the testis containing spermatocytes. (E) 104h PHS. Segmented line: CycB-positive spermatocytes. (F) Western blots probed with anti-GFP or anti-HA of proteins immunoprecipitated with anti-GFP from testis extracts of flies expressing eYFP-Fest and Rbp4-HA. Samples were dissected at 72h or 104h PHS as indicated, and all samples had RNAse A added. Negative control: Rbp4-HA alone. (G) Western blots probed with anti-GFP or anti-HA of proteins immunoprecipitated with anti-GFP from testis extracts of flies expressing eYFP-Fest and HA-Lut. Samples were dissected at 72h and 104h PHS, and all samples had RNAse A added. Negative control: HA-Lut alone.

Lut, Fest, and Rbp4 still co-immunoprecipitated in mature spermatocytes. In the hs-Bam time-course, CycB protein was not detected by immunostaining in spermatocytes at 72h PHS (Fig. 2D, curved-line bracket) but was detected in spermatocytes at 104h PHS (Fig. 2E, segmented line), just before entry into the first meiotic division. The translation of *cycB* in mature spermatocytes is not likely to be due to dissociation of either Rbp4 or Lut from Fest, as immunoprecipitation of eYFP-Fest brought down Rbp4-HA and HA-Lut at both 72h and 104h PHS (Fig. 2F,G).

### Lut is required for translational repression of *cycB* in an 8-hour window just before spermatocytes are fully mature

Loss of function of Rbp4 and Lut had different effects on the timing of CycB protein expression. Testes from either wild type, *rbp4*, or *lut* mutants were collected at 54h, 84h, 94h, and 102h PHS in the hs-Bam time-course and immunostained for CycB. In wild-type time-course testes, CycB protein was not detected in spermatocytes at the first three timepoints but was detected in spermatocytes at 102h (Fig. 3A-D). In *lut* time-course testes, CycB was detected at 94h and 102h but not at 54h or 84h PHS (Fig. 3E-H). Notably, the time at which Lut function was required to repress *cycB* translation was much later than the appearance of the Lut protein (48h PHS, Fig. 1D). In *rbp4* time-course testes, in contrast, expression of CycB protein was detected at all four time points (Fig. 3I-L).

**Figure 3.**
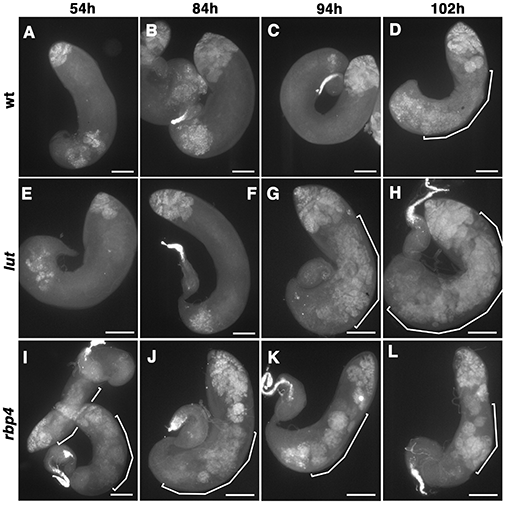
Ectopic expression of CycB occurs later in *lut* than it does in *rbp4*. (A-L) Testes from the hs-Bam time-course stained with anti-CycB. (A-D) wild-type (E-H) *lut* (I-L) *rbp4*. Segmented lines: CycB-positive spermatocytes. Scale bars 100 μm.

### Spermatocytes in *lut* or *rbp4* mutants advance to and exit the meiotic divisions six to eight hours earlier than in wild-type

Germ cells in *lut* or *rbp4* mutant males entered and exited the meiotic divisions earlier than their wild-type counterparts. In live squashed preparations viewed by phase-contrast microscopy, wild-type mature spermatocytes have a boxy nucleus and a phase-dark, round nucleolus (Fig. 4A), while spermatocytes just entering the first meiotic division have a rounder nucleus and a shrunken, crumbly nucleolus (Fig. 4B). Dividing cells are recognizable by their phase-dark array of mitochondria associated with the meiotic spindle (Fig. 4C), and post-meiotic round spermatids have a phase-light round nucleus paired with a phase-dark round mitochondrial derivative (Fig. 4D). Testes from wild-type, *lut*, or *rbp4* males in the hs-Bam time-course background were scored for the presence of germline cysts containing cells that had advanced at least as far as meiotic entry (4B) (Fig. 4E), as well as for the presence of cysts of round spermatids (4D) (Fig. 4F). At 106-108h PHS, more than half of wild-type time-course testes contained at least one cyst showing meiotic entry, whereas *lut* and *rbp4* mutants hit that benchmark at 98-100h PHS and 100-102h, respectively. Appearance of cysts of round spermatids ran about 6 hours earlier in both *lut* and *rbp4* compared to wild-type testes, as about one-third of testes from those two mutant genotypes contained at least one cyst with round spermatids at 104-106h PHS (vs. 110-112h PHS in wild type).

**Figure 4.**
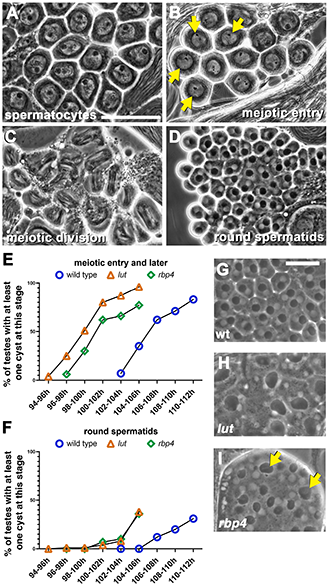
*lut* and *rbp4* spermatocytes enter and exit the meiotic divisions earlier than in wild type. (A-D) Phase-contrast imaging of unfixed wild-type testis squashes. (A) Mature spermatocytes (B) Spermatocytes initiating meiotic entry, showing rounded nuclei (arrows) and crumbly nucleoli (C) Cells in the first meiotic division (D) Round spermatids, each with a small phase-light nucleus and phase-dark mitochondrial derivative. Scale bar in (A): 50 μm, also applies to panels A-D. (E,F) Graphs showing percentage of testes from wt, *lut*, and *rbp4* with (E) at least one cyst at meiotic entry or later (i.e. including stages shown in B-D) or (F) at least one cyst with round spermatids. N=100 for each timepoint/genotype combination. (G-I) Phase-contrast imaging of testis squashes with round spermatids from (G) wild type (H) *lut* and (I) *rbp4*. Arrows in (I): larger-than-normal nebenkern, indicating at least one failed meiotic cytokinesis. Scale bar: 25 μm (in G), also applies to H and I.

Loss of function of *lut* resulted in defects in cytokinesis after meiosis I and meiosis II. In wild type, the meiotic divisions produce round spermatids with one nucleus and one phase-dark mitochondrial derivative with these two structures normally of uniform size cell-to-cell within a given cyst (Fig. 4G). In *lut* mutants, however, round spermatids typically had four nuclei and one large mitochondrial derivative per cell (Fig. 4H), suggesting complete failure of cytokinesis after meiosis I and II. In contrast, the *rbp4* mutant showed evidence of much milder cytokinesis defects (Fig. 4I, arrows).

### Spermatocyte-specific isoforms of Syp are required for CycB protein expression in mature spermatocytes

Spermatocytes express novel isoforms of *Syp*, the fly homolog of human hnRNP Q / SYNCRIP. The *syp* locus is complex, with four known promoters, a set of common/shared exons, and four possible C-terminal options, generated by alternative splicing (Fig. 5A) and encoding a variety of predicted proteins with different C-terminal regions (Fig. S1E). Results from RNA-seq, CAGE, and RT-PCR analysis indicated that promoters 1 and 4, which produce transcripts with alternate N-terminal coding sequences, fire early in spermatocyte development (Lu et al., 2020 and this study). RT-PCR confirmed that of the four annotated promoters in the *syp* locus, two (promoters 1 and 4 in Fig. 5A) were highly expressed in testis but much less so in head (Fig. S1A, first and fourth rows). Expression from promoters 1 and 4 was not detected in *bam* testes (0h PHS), which have spermatogonia but not spermatocytes, but was detected at 24h and increased by 48h PHS (Fig. S1A, first and fourth rows). Promoters 2 and 3 were expressed in testis and head and did not show stage-dependence in the time-course (Fig. S1A, second and third rows).

**Figure 5.**
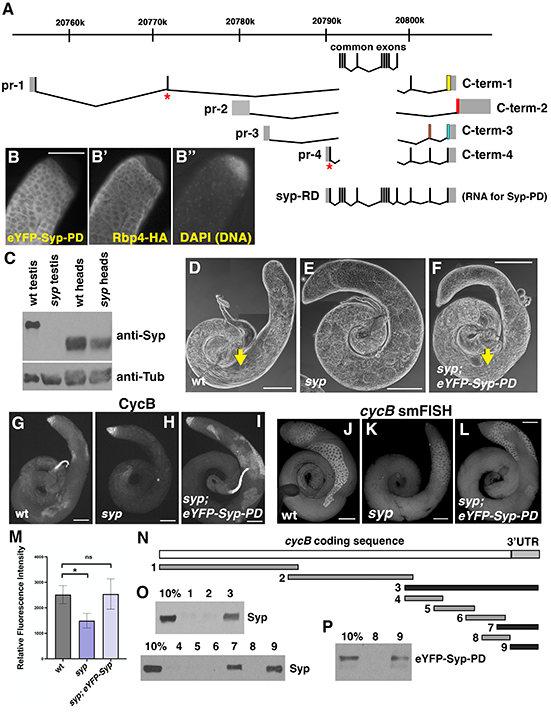
Loss of function of isoforms of Syp upregulated in testis leads to loss of CycB expression in spermatocytes. (A) The *syp* genomic locus, showing four promoters and four possible C-terminal ends, based on the FlyBase annotation. Different colors of the C-terminal exons indicate differences in protein isoforms as in Figure S1E, due to different C-terminal splice forms and/or shifts in reading frame in shared exons. The exons common to all *syp* transcripts are shown above. Asterisks indicate location of the two CRISPR-induced microdeletions that together make the *syp^dub^* allele. The complete *syp-RD* transcript is shown in full. (B-B’’) Apical tip of a testis carrying eYFP-Syp-PD and Rbp4-HA transgenes, immunostained with (B) anti-GFP and (B’) anti-HA. (B’’) DAPI (DNA). Scale bar: 100 μm. (C) Western blots probed with anti-Syp (top) and anti-Tubulin (bottom) of lysate from wild-type testes, *syp* testes, wild-type heads, and *syp* heads The *syp* mutant here and throughout the paper is *syp^dub^/Df*. (D-F) phase-contrast images of (D) wild-type, (E) *syp*, and (F) *syp; eYFP-Syp-PD* testes. Arrows indicate elongating spermatids. Scale bar: 200 μm. (G-I) anti-CycB on (G) wild type, (H) *syp*, and (1) *syp; eYFP-Syp-PD*. Scale bar: 100 μm. (J-L) smFISH with probes against *cycB*, on (J) wild type, (K) *syp*, and (L) *syp; eYFP-Syp-PD*. Scale bar: 100 μm. (M) Quantification of smFISH signal in wt, *syp*, and *syp*; *eYFP-Syp-PD* spermatocytes. Values represent the average signal measurement for three testes per genotype. Brackets denote standard deviation. Wild type vs. *syp*, p = 0.0190; wild type vs. *syp*; *eYFP-Syp-PD*, p = 0.9526 (not significant). (N) diagram of numbered biotin probes tiled across the *cycB* coding sequence and 3’UTR. Black: probes that brought down Syp; gray: probes that did not. (O) Western blots probed with anti-Syp of proteins isolated by biotin pulldown from wild-type testis extract, with probe number indicated. (P) Western blot probed with anti-GFP of biotin pulldowns with probes 8 and 9, from testis extract from eYFP-Syp-expressing flies.

Of the four 3’ splice options at the *syp* locus, two (C-term-1 and C-term-4) showed robust expression starting soon after the switch from spermatogonia to spermatocytes in the hs-Bam time-course. These two C-terminal options were not detected in head or *bam* mutant testes but appeared in testis at 24h and increased by 48h PHS in the hs-Bam time-course (Fig. S1A, fifth and eighth rows). In contrast, C-term-2 and C-term-3 showed broad expression in testis and head (Fig. S1A, sixth and seventh rows).

The predominant RNA products in testis were those in which the testis-enriched promoters were paired with the testis-enriched C-terminal splice forms, and the generally-expressed promoters were paired with the generally-expressed splice forms. RT-PCR with pairwise combinations of forward primers corresponding to either promoter 1, 2, or 4 and reverse primers from C-term-1, 2, or 4 showed PCR products from testis extracts for promoter 1 with C-term-1 and −4 only, for promoter-2 with C-term-2 only, and for promoter 4 with all three 3’ splice options (although C-term-2 was the least abundant) (Fig. S1B). These data suggest that the generally-expressed promoter 2 either (a) fires at very low levels in spermatocytes or (b) fires in a different cell type within the testis, such as spermatogonia or somatic cells.

A tagged minigene transgene termed eYFP-Syp-PD (Flybase annotation), containing a cDNA representing transcript *syp-RD* (promoter 4 + C-term-4, Fig. 5A) and 573 bp of sequence directly upstream of the predicted TSS for promoter 4, drove expression of the eYFP-Syp protein starting in early spermatocytes (Fig. 5B), at about the same time as expression of Rbp4-HA (Fig. 5B’) and broadly corresponding with the early onset of expression from promoter 4. The protein isoform encoded by the eYFP-Syp-PD transgene was entirely cytoplasmic, consistent with the lack of a predicted NLS (Fig. S1E). Reporter transgenes made with either promoter 1 or promoter 4 and with C-term-1 showed nuclear expression in early spermatocytes (consistent with the presence of a predicted NLS encoded by C-term-1), and both nuclear and cytoplasmic localization later (Fig. S1C,D).

Co-injection of CRISPR RNAs targeting the unique N-terminal coding sequences transcribed from promoters 1 and 4 resulted in creation of a *syp* allele (*syp^dub^*) with a pair of microdeletions, one downstream of each promoter, that resulted in frameshifts and early termination of the corresponding proteins (Materials and Methods). Flies that were *syp^dub^/Df* (hereafter referred to as *syp*) were viable and female-fertile but male-sterile. Syp protein was not detected by anti-Syp (guinea pig) western blot in testes from *syp* flies (Fig. 5C). Curiously, heads from *syp* flies showed a reduction in Syp protein. Expression of Syp protein in somatic cells was highlighted by immunostaining with anti-Syp (rabbit) when germline Syp was absent in *syp* testes (Fig. S1G vs. F), and co-staining for Traffic Jam (Tj) (Fig. S1H, H’) indicated that the isoforms of Syp present in somatic cyst cells can localize to the nucleus (arrows) as well as the cytoplasm. These data suggest that *syp* RNA isoforms detectable in testis and originating from promoters 2 or 3 are expressed in somatic cells.

Syp is needed in males for progression through the meiotic divisions and post-meiotic differentiation. In wild-type testes, elongating spermatids were clearly visible (Fig. 5D, arrow), whereas in *syp* mutant testes, no elongating spermatids were observed, and arrested spermatocytes accumulated (Fig. 5E). The eYFP-Syp-PD transgene partially rescued the *syp* phenotype, with elongating spermatids visible (Fig. 5F, arrow).

Syp function is required for accumulation of CycB in mature spermatocytes. In *syp* mutant testes, CycB protein was expressed normally in spermatogonia but did not reappear in mature spermatocytes (Fig. 5, H vs. G). Single molecule FISH (smFISH) with probes against *cycB* indicated that the *cycB* RNA was still present in *syp* mutant spermatocytes, although at lower levels than in wild type (Fig. 5, K vs. J; quantified in M). One copy of the eYFP-Syp-PD transgene was sufficient to rescue expression of CycB protein in mature spermatocytes in a *syp* mutant background (Fig. 5I) as well as restore *cycB* RNA levels in spermatocytes (Fig. 5L, quantified in M). The ability of the exclusively-cytoplasmic eYFP-Syp-PD to rescue CycB protein expression argues that cytoplasmic rather than nuclear functions of Syp are important for CycB protein to be expressed in mature spermatocytes.

Syp protein binds to the 130 nt *cycB* 3’UTR expressed in spermatocytes, suggesting direct action. *In vitro* transcribed biotin-labeled probes corresponding to portions of the *cycB* RNA (diagrammed in Fig. 5N) were incubated with testis extracts from wild-type flies and captured with streptavidin beads. Associated proteins were then analyzed by anti-Syp western blot (Fig. 5O). Syp protein associated with probes containing the short spermatocyte 130 nt *cycB* 3’UTR but not with probes representing the *cycB* protein coding sequences. The eYFP-Syp-PD reporter protein also bound the *cycB* 3’UTR, in similar biotin pulldown assays from testes from males carrying the eYFP-Syp-PD transgene (Fig. 5P).

The *cycB* RNA was detected, although at levels considerably lower than wild type, in late-stage and mature spermatocytes in *syp* mutants, past the stage at which *cycB* is normally translated in wild type. In testes from the hs-Bam time-course, smFISH revealed the presence of *cycB* RNA in wild-type mature spermatocytes from 100h through 112h PHS (Fig. 6A-D), with *cycB* RNA levels dropping in most cysts by 116h (Fig. 6E). In *syp* mutant spermatocytes, *cycB* RNA was detected at lower levels than wild type throughout the first three timepoints (Fig. 6F-H), with a drop in *cycB* levels in some cysts at 112h (Fig. 6I) and in most cysts by 116h (Fig. 6J).

**Figure 6.**
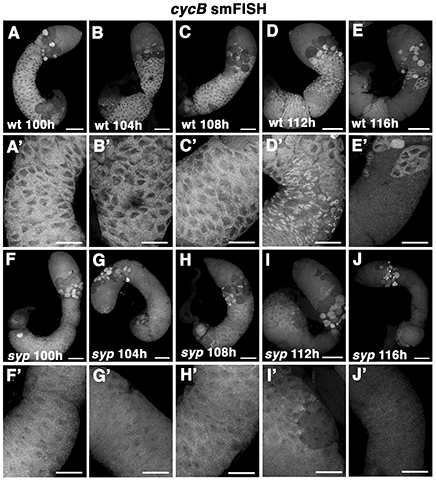
*cycB* RNA levels are lower but detectable through 108h PHS in *syp* mutants. (A-J) Testes from timepoints in the hs-Bam time-course probed for *cycB* RNA by smFISH. (A-E) wild-type time-course testes. (A’-E’) higher magnification of spermatocyte regions of (A-E). (F-J) *syp* mutant time-course testes. (F’-J’) higher magnification of spermatocyte regions of (F-J). 100h, 104h, 108h, 112h, and 116h PHS as indicated for both genotypes. Scale bars in A-E and F-J: 100 μm. Scale bars in A’-E’ and F’-J’: 50 μm.

### Spermatocytes mutant for *syp* arrest before pro-metaphase

Wild-type function of Syp in spermatocytes is required for successful entry into and progression through the meiotic divisions. Wild-type time-course spermatocytes at 72h PHS showed partially condensed bivalents appearing as chunky crescents near the nuclear membrane (Fig. 7A’, white arrows). By 110h PHS most had reached a stage at which nucleoli had shrunk or disappeared and the chromosomes had condensed (Fig. 7B’, white arrows) in advance of congressing to the center of the nucleus for metaphase (not shown). Spermatocytes mutant for *syp*, in contrast, showed normal chunky crescent-shaped chromosomes at 72h (Fig. 7C’) but show only partially-condensed chromosomes at 110h (Fig. 7D’, arrows) and did not break down their nucleoli (Fig. 7D, arrows). Metaphase figures (chromosomes in a tight knot at the center of the nucleus) were never observed in *syp* mutants. Unlike *twine* and *boule* mutants, which skip the meiotic divisions and complete a convincing (*twine*) or modest (*boule*) amount of post-meiotic differentiation (Alphey et al., 1992; Eberhart et al., 1996), *syp* germ cells did not show any signs of post-meiotic differentiation, such as formation of the nebenkern which normally takes place in round spermatids (as seen in wild type in Fig. 4D).

**Figure 7.**
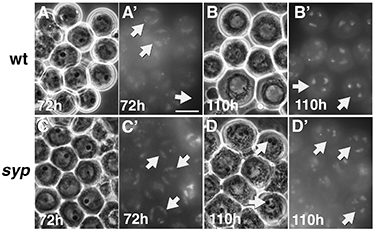
Spermatocytes mutant for *syp* arrest before prometaphase. Phase-Hoechst images of spermatocytes in squashed preparations of testes from the hs-Bam time-course on (A-B’) wild-type and (C-D’) *syp* at 72h or 110h PHS as indicated. (A, B, C, and D) Phase-contrast. (A’, B’, C’, and D’) Hoechst (DNA). Arrows in A’ and C’: cells with chunky, crescent-shaped chromosomes; arrows in B’: cells with condensed chromosomes. Arrows in D: cells with intact nucleoli. Arrows in D’: cells with partially condensed chromosomes. Scale bar in A’ is 25 μm and applies to all panels.

### Syp binds to Fest and interacts with Rbp4 and Lut dependent on Fest

The cytoplasmic isoform of the Syp protein encoded by Syp-PD co-immunoprecipitated Fest, in an interaction that did not require Rbp4. Immunoprecipitation of eYFP-Syp with anti-GFP from testis extracts brought down HA-Fest (Fig. 8A). This interaction remained strong in the *rbp4* mutant background, suggesting that Fest and Syp do not require Rbp4 for their interaction. The translation of *cycB* in mature spermatocytes is not likely due to dissociation of Syp from Fest, as immunoprecipitation of eYFP-Syp brought down HA-Fest at both 72h and 104h PHS (Fig. 8B).

**Figure 8.**
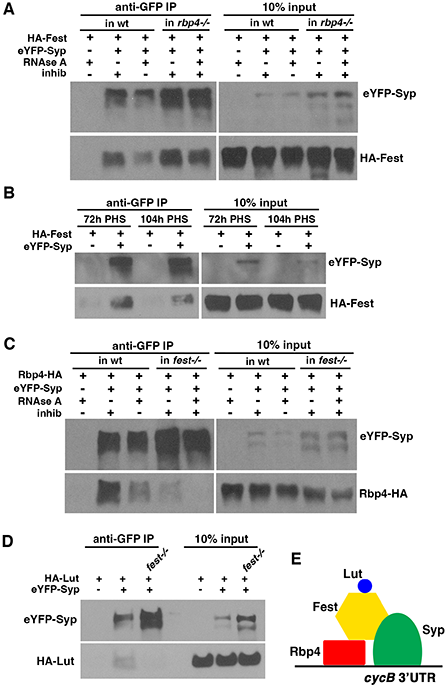
Syp interacts with Fest and co-immunoprecipitates with Rbp4 and Lut in the presence of Fest. (A) Western blots probed with anti-GFP or anti-HA of anti-GFP immunoprecipitates from testes expressing eYFP-Syp and HA-Fest, with either RNAse A or RNAse inhibitor, as indicated. Negative control: HA-Fest alone. Flies were either wt (first three lanes) or mutant for *rbp4* (lanes 4 and 5). (B) Western blots probed with anti-GFP or anti-HA of anti-GFP immunoprecipitates from hs-Bam time-course testes expressing eYFP-Syp and HA-Fest, collected at 72h and 104h PHS, as indicated. Negative controls: HA-Fest only. (C) Western blots probed with anti-GFP or anti-HA of anti-GFP immunoprecipitations from testes expressing eYFP-Syp and Rbp4-HA, with either RNAse A or RNAse inhibitor, as indicated. Negative control: Rbp4-HA alone. Flies were wt (first three lanes) or mutant for *fest* (lanes 4 and 5). (D) Western blots probed with anti-GFP or anti-HA of anti-GFP immunoprecipitates from testes also expressing eYFP-Syp and HA-Lut. All samples included RNAse A. Flies were wt or mutant for *fest*, as indicated. Negative control: HA-Lut alone. (E) Schematic of proposed complex, with Syp and Rbp4 both able to bind the *cycB* 3’UTR, and Fest acting as a scaffold protein further connecting Rbp4, Syp, and Lut in an RNA-independent manner.

Rbp4-HA also co-immunoprecipitated with eYFP-Syp, but this interaction required Fest. Immunoprecipitation with anti-GFP from testes expressing both eYFP-Syp and Rbp4-HA brought down Rbp4-HA (Fig. 8C), but this interaction was reduced in the presence of RNAse and in a *fest* mutant background (Fig. 8C), suggesting that the physical interaction between Rbp4 and Syp may be mediated by their individual interactions with Fest and with RNA. Similarly, HA-Lut co-immunoprecipitated with eYFP-Syp in the presence but not absence of Fest (Fig. 8D), in the presence of RNAse. Together these data suggested that in addition to being bound to the *cycB* 3’UTR, Syp also associates with the Rbp4-Fest-Lut complex via its binding with Fest (Fig. 8E).

### In the absence of Rbp4, Syp is not required for *cycB* translation in mid-stage spermatocytes

Analysis of CycB protein expression in testes from *rbp4 syp* double-mutant flies revealed that Syp was not required for CycB protein expression in mid-stage spermatocytes in the absence of Rbp4. In the *rbp4 syp* double mutant, a puff of CycB protein expression was detected early (Fig. 9F), arguing against the possibility that Rbp4 acts upstream to block *cycB* translation by Syp in mid-stage spermatocytes. In *lut syp* double mutants, in contrast, CycB was not detected in spermatocytes (Fig. 9E) – comparable to testes mutant for *syp* alone (Fig. 9D). The result that Syp is still required for CycB expression in late-stage spermatocytes in the absence of Lut indicates that Syp does not promote *cycB* translation by blocking repressive action by Lut.

**Figure 9.**
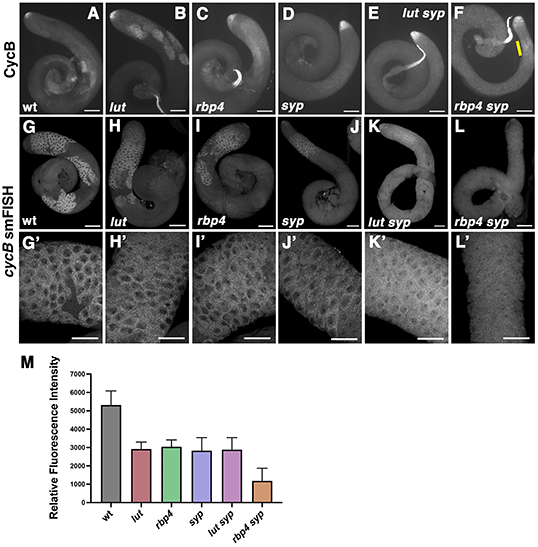
In the absence of Rbp4, Syp is not required for *cycB* translation in mid-stage spermatocytes. (A-F) Immunofluorescence images of whole testes stained with anti-CycB. (A) wild type (B) *lut* (C) *rbp4* (D) *syp* (E) *lut syp* (F) *rbp4 syp*. Yellow bar in (F) highlights a puff of CycB expression. Scale bar: 100 μm. (G-L) Whole testes probed for *cycB* RNA by smFISH. (G) wild type (H) *lut* (I) *rbp4* (J) *syp* (K) *lut syp* (L) *rbp4 syp*. Scale bar: 100 μm. (G’-L’) high magnification views of spermatocyte regions in (G-H). Scale bar in (G’-L’) is 50 μm. (M) Quantification of smFISH signal in wt, *lut*, *rbp4*, *syp*, *lut syp*, and *rbp4 syp* spermatocyte regions. Values represent the average signal measurement for three testes per genotype. Brackets denote standard deviation. P values, vs. wild type: *lut*, p = 0.0181; *rbp4*, p = 0.0209; *syp*, p = 0.0149; *lut syp*, p = 0.0148; *rbp4 syp*, p = 0.0024.

smFISH with *cycB* probes revealed that *cycB* RNA levels were lower in the *lut*, *rbp4*, and *syp* single mutants and the *lut syp* double mutant than in wild type (Fig. 9H-K vs. G; quantified in M) and even lower in the *rbp4 syp* double mutant (Fig. 9L, quantified in M), suggesting that Rbp4 and Syp may act In parallel to promote stability of the *cycB* RNA. The fact that CycB protein is detected early in spermatocyte development in *rbp4 syp* but not later (as in *rbp4* alone) could be due to a continued decrease in *cycB* RNA abundance as spermatocytes mature and/or changes in the translational environment in later spermatocyte stages (see Discussion).

## DISCUSSION

Our work reveals that during meiotic prophase in *Drosophila* the male germline developmental program controls timing of expression of the core G2 cell cycle regulator CycB by a complex of cell-type-specific translational repressors interacting with a partner that stabilizes target RNAs and may also activate their translation. Three of the earliest genes newly expressed when *Drosophila* spermatocytes enter meiotic prophase encode proteins that interact in a complex that is recruited to the short 130 nt 3’UTR of the *cycB* RNA once it becomes transcribed in spermatocytes. The Rbp4 protein is required to block translation of the *cycB* RNA in mid-stage spermatocytes while function of Lut is required to block translation of *cycB* in late-stage, not-quite-mature spermatocytes. Thus, when *cycB* transcription resumes from the spermatocyte-specific promoter, the newly-expressed transcripts enter a cytoplasm already prepared for their translational repression. Complementing this cell-type-specific repression of translation, spermatocytes also express a cytoplasmically localized protein isoform of the hnRNP R/Q homolog Syp, which binds strongly to the *cycB* 3’UTR and Fest and is required for the normal ability of mature spermatocytes to express CycB protein just prior to entry into the meiotic divisions.

Loss of function of either Lut or Rbp4 resulted in premature entry into the meiotic divisions and early progression to haploid round spermatids, with a difference of about 6-8 hours from wild type controls, indicating that early expression of CycB can be permissive for early meiotic entry but not sufficient to drive it. If it were sufficient, CycB accumulation at 54h PHS in the *rbp4* mutant would result in meiotic entry much earlier than the observed 100-102h PHS timepoint. It is likely that spermatocytes require additional critical core G2 cell cycle regulators to undergo the G2/M transition of meiosis I. If such additional regulators only become active by the stage equivalent to 90-96h PHS, premature expression of CycB may not be sufficient to drive the G2/M transition prior to that time.

The difference in timing of CycB protein expression in *rbp4* vs. *lut* points to two distinct states in the regulation of *cycB* translation. Rbp4 is required for repression of *cycB* translation in mid-stage spermatocytes (54h-94h PHS), while Lut function is not required to prevent CycB protein accumulation at this stage. In late-stage spermatocytes (94h-102h PHS), in contrast, function of Lut is required for translational repression of *cycB*, and, <u>in the absence of Lut</u>, Rbp4 function is either off or not sufficient to repress *cycB* translation.

Rbp4, Lut, and Syp participate in a multi-protein complex anchored by Fest. As we have yet to detect changes in the composition of the complex in early vs. late spermatocytes, it remains a mystery as to how translational repression by the complex is eventually relieved in mature spermatocytes preparing for the G2/M transition of meiosis I. However, using results from the epistasis tests (Fig. 9), we can postulate possible regulatory relationships among the interacting proteins in the complex bound to the *cycB* 3’UTR in mid-stage, late-stage, and mature spermatocytes. The three models presented here are not meant to rule out other possibilities, but each one is compatible with the data from single mutants, double mutants, and our time-course data. In particular, expression of CycB protein in young *rbp4 syp* spermatocytes indicates that Rbp4 does not act upstream of Syp (for example, by blocking *cycB* translation by inhibiting action of Syp). Likewise, lack of detectable CycB protein in *lut syp* spermatocytes would suggest that Syp does not act upstream of Lut (for example, allowing translation by turning off repression by Lut).

The first model (Fig. 10A) postulates that (a) Rbp4 represses translation of *cycB* in mid-stage spermatocytes (^~^54-94h PHS), subject to reversal by factor Y; (b) Lut represses translation of *cycB* in late-stage spermatocytes (^~^94-102h PHS), subject to reversal by factor X in mature spermatocytes (^~^102h+ PHS); and (c) Syp functions primarily to stabilize *cycB* RNA and maintain it at sufficiently high levels to produce detectable amounts of CycB protein in mature spermatocytes. Levels of *cycB* RNA are clearly reduced in the absence of Syp (Figs 5, 6, and 9). One possible explanation is that occupancy of the short *cycB* 3’UTR by Syp protein blocks other RNA-binding proteins that bind 3’UTRs and recruit the deadenylase complex to target RNAs (Fukushima et al., 2019; Arvola et al., 2020; Semotok et al., 2005). As noted above, *cycB* RNA was detected at 104 and 108h in the *syp* mutant (Fig. 6), albeit at lower than normal levels. However, the baseline translational environment at that stage may be less permissive than the environment in mid-stage spermatocytes, such as those that express CycB protein in the *rbp4 syp* double mutant (Fig. 9).

**Figure 10.**
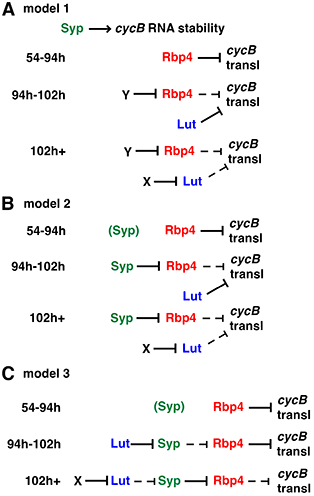
Three models for Lut, Syp, and Rbp4 function in regulating *cycB* translation and stability. Solid lines indicate active promotion or repression; dotted lines indicate abrogated function. Each protein is color-coded to match the diagram in Figure 8. Not shown, but contributing to all three models: Syp and Rbp4 act in parallel to stabilize the *cycB* RNA. See Discussion.

The second model (Fig. 10B) presents the possibility that Syp could be factor Y, suppressing Rbp4 function in maturing spermatocytes, while still acting in parallel with Rbp4 to stabilize the *cycB* RNA. In this model, Syp is present in mid-stage spermatocytes but not capable of blocking translational repression by Rbp4 until late-stage spermatocytes, setting up the requirement for Lut in the 94-102h window as the last bastion of *cycB* translational repression, until Lut function itself is counteracted at 102h by factor X.

In the third model (Fig. 10C), Lut, Syp, and Rbp4 function in a sequential repressive cascade. In mid-stage spermatocytes, as in model 2, Rbp4 represses *cycB* translation, with Syp present in the complex but not capable of interfering with Rbp4 function. In late-stage spermatocytes, Syp is capable of blocking Rbp4 repressive activity, but is prevented from doing so by Lut. Thus, loss of function of Lut allows *cycB* translation in this late-stage window. In mature spermatocytes, factor X counteracts Lut function, releasing Syp to antagonize Rbp4 function, thus allowing *cycB* translation.

Consistent with these models, Syp and mammalian Syp homologs have been implicated in regulation of RNA stability, localization, and translation. In *Drosophila*, Syp has been shown to regulate developmental and cell fate decisions in a diverse set of cell types and tissues, with specific RNA targets identified in some cases. Syp functions in the oocyte to promote dorso-anterior localization of the *gurken* RNA, binding the *gurken* localization signal (McDermott et al., 2012). In the larval neuromuscular junction, Syp acts post-synaptically to block synaptic overgrowth, binding multiple target RNAs (McDermott et al 2014). Syp binds the *MSP-300* RNA, co-localizes with it *in vivo* to ribosome-dense granules (Titlow et al., 2020), and promotes translation of *MSP-300* in a neural-activity dependent manner. In the *Drosophila* mushroom body, Syp and another RNA-binding protein Imp appear in opposing temporal gradients and regulate cell fate decisions (γ, α’/β’, or α/β neurons), with Syp antagonizing and Imp promoting expression of Chinmo protein without either factor affecting *chinmo* RNA abundance (Liu et al., 2015). Syp promotes neuroblast decommissioning by upregulating expression of the Prospero protein (Yang et al., 2017) via stabilization of the long form of the *prospero* RNA expressed in larvae (Samuels et al., 2020). From this extensive literature, it is clear that Syp can stabilize RNAs and have repressive or activating effects on translation of specific target RNAs, sometimes even in a single cell type (McDermott et al., 2014).

## MATERIALS AND METHODS

### Fly genetics and husbandry

Flies were raised on dextrose/cornmeal food at 20°C for stocks and 25°C for crosses. The *rbp4* mutant flies were *rbp4^LL06910^/Df(3R)Exel6169*. *Df(3R)Exel6169* is from Bloomington stock #7648, and the *rbp4^LL06910^* allele was from the Drosophila Genomics Resource Center (DGRC, #141934). The *lut^1^* and *syp^dub^* alleles were generated by CRISPR as described below. The *lut* mutant flies were *lut^1^/Df(2R)BSC131*. *Df(2R)BSC131* is from Bloomington stock #9296. The *syp* mutant flies were *syp^dub^/Df(3R)BSC141*, with *Df(3R)BSC141* from Bloomington line #9501. For the hs-Bam time-course, ^~^ 40 *w*; *hs:Bam-HA*/*CyO*; *bam^Δ86^*/*TM3* virgin females were mated to *w*;; *bam^1^*/*TM6B* males for wild-type controls. For *lut* in the time-course: *w*; *hs:Bam-HA*, *lut^1^*/*CyO*; *bam^Δ86^*/*TM3* x *w*; *Df(2R)BSC131*; *bam^1^*/*TM6B*. For *rbp4* in the time-course: *w*; *hs:Bam-HA*/*CyO*; *Df(3R)Exel6169*, *bam^Δ86^*/*TM3* x *w*;; *rbp4^LL06910^*, *bam^1^*/*TM6B*. For *syp* in the time-course: *w*; *hs:Bam-HA*/*CyO*; *Df(3R)BSC141*, *bam^Δ86^*/*TM3* x *w*;; *syp^dub^*, *bam^1^*/*TM6B*.

### Hs-Bam time-course

For the hs-Bam time-course, fly bottles (plastic) replete with mid-stage and late-stage pupae were weighted down and placed in a 37°C water bath with water nearly up to the lip of the bottle. After 30 minutes the bottles were cleared of any adults that may have eclosed, labeled with date and time of heat shock, and returned to 25°C for the designated time post heat shock. The clock starts on hours-post-heat-shock at the end of the 30-minute 37°C incubation. Note that time-course testes regain a population of *bam* mutant spermatogonia at the apical end of the testis during the incubation at 25°C after heat shock, due to mitotic proliferation of *bam* mutant early germ cells that avoided forced differentiation.

### CRISPR alleles

Microdeletions were generated by CRISPR using the technique described in Bassett et al., 2013. Genomic target sites were chosen early enough in the coding sequence to cause a significant disruption to the encoded protein, and containing the sequence GN[N_18_]NGG in either direction. This sequence was then incorporated into the middle of a forward primer after changing the first N to G, and removing NGG, to produce the following oligo: GAAATTAATACGACTCACTATA**GG(N_18_)**GTTTTAGAGCTAGAAATAGC, containing a T7 promoter (underlined) followed by the adjusted gene-specific sequences (bold). Template-less PCR was performed with this primer and a common reverse primer (AAAAGCACCGACTCGGTGCCACTTTTTCAAGTTGATAACGGACTAGCCTTATTTTAACTTGCTATTTCTA GCTCTAAAAC) as described, then cleaned up using a PCR purification kit (Bassett et al., 2013). 250 ng of PCR product were used as a template for an *in vitro* transcription reaction (MEGAscript T7 kit), incubated at 37°C for 2 hours. After treatment with DNAse for 15 minutes, the resulting RNA was purified by Trizol prep. For each target, the guide RNA was then injected into Act5-Cas9 embryos at a concentration of 100 ng/μl. Surviving adults were individually crossed to the deficiency line uncovering the gene of interest, and the resulting progeny were screened for the phenotypes observed previously via testis-specific RNAi knockdown (VDRC #33011 for *syp* and VDRC #29664 for *lut*). The gene-specific GN[N_18_]NGG sequence for *lut* was GGGCCAGCAGGGATTCCATTTGG and the resulting microdeletion in *lut^1^* was GGGCCAGCAGGG**xxxxx**ATTTGG. The mutated coding sequence encodes only the first 45 amino acids of Lut (out of 132), three additional aberrant residues, and a stop codon. For *syp*, the gene-specific sections of the guide RNAs were GTACCCGTTATCAAGCCCATTGG (downstream of promoter 1) and GGAAGCCTCCAAAGTGCAGAAGG (downstream of promoter 4), and their co-injection generated the allele *syp^dub^* allele, which contains two microdeletions in tandem: GTACCCGTTATCAAGC**x**CATTGG (pr1) and GGAAGCCTCCAAA**xxxxxxxx**GG (pr4). The mutated coding sequence downstream of promoter 1 encodes the first 26 correct amino acids, 82 aberrant residues, and a stop codon, while the mutated coding sequence downstream of promoter 4 encodes the first 9 correct amino acids, one aberrant residue, and a stop codon.

### Transgenes

The transgenes described below are diagrammed in Fig. S2. The eYFP-Syp-PD reporter, built in pBlueScript-KS and then moved into the *Not*I/*Spe*I sites of pCaSpeR4, consisted of: *Not*I – *syp* promoter-4 (573 bp, directly 5’ of FlyBase-annotated *syp-RD* start site) and the *syp-RD* 5’UTR (223 bp) – *Xba*I/*Spe*I [non-recleavable] – *eYFP* coding sequence (717 bp) – *Xba*I – *syp-RD* coding sequence (2124 bp) – *Xba*I – *syp-RD* 3’UTR and 3’ downstream genomic sequence (811 bp combined) – *Spe*I. The eYFP-Syp-PT and eYFP-Syp-PV reporters were built in pBS-KS and moved into the *Not*I/*Spe*I sites of pCaSpeR4 to which the SV40 terminator had been added. Note that the promoter for *syp-RD* and *syp-RV* is the same, while the C-terminal coding sequence and 3’UTR are identical for *syp-PV* and *syp-PT*. The eYFP-Syp-PV transgene consisted of: *Not*I – *syp* promoter-4 (573 bp, directly 5’ of annotated *syp-RD/RV* start site) and the *syp-RD/RV* 5’UTR (223 bp) – *Xba*I/*Spe*I [non-recleavable] – *eYFP* coding sequence (717 bp) – *Xba*I – *syp-RV* coding sequence (2037 bp) – *Xba*I – *syp-RT/RV* 3’UTR and 3’ downstream genomic sequence (804 bp combined) – *Spe*I. The eYFP-Syp-PT transgene consisted of: *Not*I – *syp* promoter-1 (494 bp, directly 5’ of annotated *syp-RT* start site) and the *syp-RT* 5’UTR (508 bp) – *Xba*I/*Spe*I [non-recleavable] – *eYFP* coding sequence (717 bp) – *Xba*I – *syp-RT* coding sequence (2013 bp) – *Xba*I – *syp-RT/RV* 3’UTR and 3’ downstream genomic sequence (804 bp combined) – *Spe*I.

The HA-Lut and V5-Lut reporters, built in pBS-KS and moved to the *Not*I/*Xho*I sites of pCaSpeR4, consisted of: *Not*I – 232 bp 5’ genomic and 199 bp 5’UTR of *lut* – *Xba*I – V5 or 3xHA coding sequence (42 bp and 126 bp, respectively) – *Eco*RI – *lut* coding sequence plus an intron (467 bp total), *lut* 3’UTR and 3’ downstream genomic sequence (534 bp combined) – *Xho*I.

HA-Fest, like eYFP-Fest (Baker et al., 2015), was built in pBS and then moved into the *Xba*I/*Xho*I sites of pCaSpeR4 to which the SV40 terminator had been added. The plasmid contained: *Xba*I – the *fest* promoter (590 bp, directly 5ʹ of the annotated transcription start site) and the *fest* 5ʹ UTR (275 bp) – *Spe*I – *3xHA* coding sequence (126 bp) – *Sma*I – the *fest* coding sequence (1542 bp) – *Eco*RI – the *fest* 3ʹ UTR and 3’ downstream genomic sequence (1255 bp total) -*Eco*RI-*Xho*I – *SV40* terminator – *Sal*I/*Xho*I [non-recleavable]. Transgenic flies were generated by microinjection either at BestGene (eYFP-Syp and HA-Fest) or in-house (the remainder).

Coding sequences for *syp* and *fest*, as well as the 5’UTR of *syp-RT*, were amplified from cDNA generated from wild-type testis RNA. All other gene-specific sequences listed above were amplified from wild-type genomic DNA.

### RT-PCR

RNA was collected from wild-type testes and heads, as well as hs-Bam time-course testes from 0, 24, and 48 hours post-heat-shock. Reverse transcription with oligo-dT primer and Ready-to-Go You-Prime First-Strand Beads (Cytiva #27926401) was performed on 1 μg of each RNA sample. PCR reactions were done in a 25 μl volume with 2 μl cDNA each. BioMix Red (Bioline #BIO-25006) was used for all reactions *except* that which amplified C-term-4, which is GC-rich and required MyTaq (Bioline #BIO-21105). The amplification ran for 30 cycles (Fig S1A) or 35 cycles (S1B). Primers for Fig. S1A: the common reverse primer for assaying promoters was 5’-GTTCTCGAATAGCGGAATCAG-3’. Forward primers: promoter 1: 5’-GTCAGAAACACTCGCATGCAAA-3’; promoter 2: 5’-ACTCACTTGGATACACAGCG-3’; promoter 3: 5’-TGTGCAACGCCGAGCAGAGT-3’; promoter 4: 5’-GATGGAAGCCTCCAAAGTGC −3’. Common forward primer (for use on C-term 1, 2, and 3): 5’-CAGGACGTCTCAGAGGATAA-3’. Forward primer for use on C-term 4: 5’-GTGAATACGACTACTTTTACGAC-3’. Reverse primers: C-term-1: 5’-TGCAGCTCCAGCATAAGGCT-3’; C-term-2: 5’-TACCGACTACGTATTCACGG-3’; C-term-3: 5’-CGACCAACTCCCGTATTCAC-3’ ; C-term-4: 5’-GCTCCAGCTGGTAAATTTTGT-3’. For the *GAPDH2* controls, forward and reverse primers were 5’-CCGTTCATGCCACCACCGCT-3’ and 5’-GCCACGTCCATCACGCCACA-3’, respectively. For Fig. S1B, the relevant isoform-specific forward and reverse primers listed above were used for promoters 1, 2, and 4 and C-terms 1 and 2, while a new primer was used as a reverse primer for C-term 4: 5’-ACGACCAACTCCCATAAGGCT-3’. BioMix Red was used for all 9 reactions shown in Fig. S1B.

### Biotin pulldowns

Probes were cloned into pBS-KS in the orientation *Spe*I 5’->3’ *Bam*HI, with the T7 promoter 5’ of the *Spe*I site. The following probes were used, with the nt numbers corresponding to the 1593 nt protein coding sequence and the short spermatocyte 3’UTR (130 nt) of *cycB-RA*, starting with the first base of the coding sequence (total, 1723 nt): #1, 1-631; #2, 589-1187; #3, 1117-1723; #4, 1117-1292; #5, 1253-1440; #6, 1392-1574; #7, 1534-1723; #8, 1463-1593; #9, 1594-1723. Plasmids containing the probe templates were linearized with *Bam*HI for several hours, and then run on a gel and purified using the Zymogen gel cleanup kit. For *in vitro* transcription, T7 polymerase and biotin labeling mix (Roche #11685597910) were used according to the package directions. After 2 hours of transcription at 37°C, 2 μl DNAse was added and reactions were returned to 37°C for an additional 30 minutes. Probes were cleaned of unincorporated nucleotides using NucAway columns (Thermo Fisher #10070). Fifty pairs of testes per sample (plus 5 pairs per genotype as input) were mechanically lysed in [100 μl/sample + 10 μl extra] lysis buffer (100 mM NaCl, 50 mM Tris, 4 mM EDTA, 1% NP40, 1 μl/ml SUPERaseIN (Thermo Fisher #AM2694), and 1x cOmplete Protease Inhibitor (Roche #11836153001)) with a 1 cc syringe and 25×5/8 needle, rocked at 4°C for 30 minutes, and centrifuged at full speed for 3 minutes at 4°C to pellet insoluble material. While the lysate was rocking, 50 μl suspended streptavidin MagneSphere beads (Promega #Z5481) per sample were washed 3×5 minutes in 0.5x SSC, with a 6-tube magnetic stand (Thermo Fisher #AM10055) used to secure beads between washes. 10% input (10 μl lysate) was removed for each genotype, supplemented with Laemmli sample buffer to 1x, and boiled; the remaining 100 μl lysate per sample was pre-cleared by incubating with just-prepped beads for 30 minutes at RT. Pre-cleared lysate was removed from those beads and transferred into fresh tubes in 100 μl aliquots (one per probe), into each of which 500 ng of a single biotin-labeled probe was added. Samples were rocked 30 minutes at RT, after which the lysate/probe mixture was incubated with fresh 50 μl streptavidin beads for 30 minutes at RT with rocking. The beads were washed 5×10 minutes in 1 mL cold lysis buffer at 4°C with rocking. Laemmli sample buffer was added to a concentration of 1x before the samples were boiled. After boiling, samples (including beads) were centrifuged briefly, and the supernatant was loaded onto a protein gel and analyzed by western blot.

### Immunoprecipitations

Protein A Dynabeads (Thermo Fisher #10002D) were used with rabbit anti-HA (Cell Signaling #3724S), and pan-mouse IgG Dynabeads (Thermo Fisher #11041) were used for anti-GFP (mouse, Millipore Sigma #11814460001) and anti-V5 (mouse, Thermo-Fisher #R960-25). The 6-tube magnetic stand from Thermo-Fisher was used, as above, to segregate the beads from liquid between washes during antibody conjugation and immunoprecipitations. 50 μl of suspended beads per sample were washed twice in 1 mL 3% BSA / PBSTw (PBS + 0.1% Tween), brought back up to the original volume (50 μl per sample) and split into two aliquots: 20 μl/sample to be used in the pre-clear step, and 30 μl/sample to be used for antibody conjugation and subsequent immunoprecipitation. To conjugate antibody to beads, excess BSA wash was removed from the suspended beads, and beads were incubated with lysis buffer and antibody in a ratio of 30 μl (suspended) beads : 200 μl lysis buffer (see below, but without protease inhibitor) : 2 μl of antibody per IP. For example, for 5 immunoprecipitation reactions: 10 μl antibody, 1 mL lysis buffer, 150 μl suspended beads. Beads were incubated with antibody for 1 hour at room temperature, then split into individual aliquots (one per IP) and washed 3× 5 minutes in 1 mL 0.2M triethanolamine pH 8.2. Beads were then treated for 30 minutes at RT with triethanolamine to which dimethylpimelimidate had been freshly added to a final concentration of 5.4 mg/mL. After a 15-minute wash in 1 mL 50 mM Tris, pH 7.5, and a final wash with 1 mL PBSTw, beads could be stored at 4°C for 1-2 days before use. On the day of the immunoprecipitation, beads were stripped of non-covalently bound antibody via two quick washes in 1 mL 100mM glycine, pH 2.5. After a wash with PBSTw, they were blocked in 1 mL 10% BSA / 50mM Tris for 1 hour before a 5-second rinse in PBSTw.

Testes (55 pairs per sample) were mechanically lysed in 220μl cold lysis buffer (135 mM NaCl, 20 mM Tris, 10 mM EDTA, 1% NP40, 10% glycerol, 1x cOmplete protease inhibitor) per sample with a 1 cc syringe and 25×5/8 needle, at 4°C. Samples were then incubated for 20 minutes on a rocker at 4°C before samples were centrifuged at full speed for 3 minutes to pellet insoluble material. The supernatant was collected and split evenly if needed (for “+ RNAse” vs. “+ inhibitor”). For “+ RNAse” samples, RNAse A (Thermo Fisher #EN0531) was added to a final concentration of 100 μg/mL; for “+ inhibitor” samples, SUPERasin was added to a final concentration of 1U/μl. To pre-clear the lysate, each sample was then transferred to a fresh tube with 20 μl suspended BSA-washed beads (from above; excess BSA wash removed just before using) and incubated at 4°C for 45 minutes with rocking. 20 μl of lysate was then removed from each sample to serve as 10% input, to which 5 μl 5x Laemmli sample buffer was added before boiling. The remaining 200 μl lysate for each sample was transferred to a fresh tube containing antibody-conjugated beads for the immunoprecipitation step (3-4 hours at 4°C, rocking). Beads were then washed 2x 10 minutes in 1 mL cold lysis buffer at 4°C while rocking. Bound proteins were eluted from the beads in 40 μl elution buffer (10 mM EDTA, 50 mM Tris pH 8, 1% SDS, and 1x cOmplete protease inhibitor) at 70°C for 30 minutes with frequent mixing. The resulting 40 μl eluate was transferred to a fresh tube, and 10 μl 5x Laemmli sample buffer was added. Samples were boiled 10 minutes and frozen until analysis by western blot.

### Western blotting

Samples were run on 10% or 4-15% TGX pre-cast gels with 10 wells with 50 μL well volume (Bio-Rad, #4561034 and #4561084). Proteins were transferred overnight onto PVDF membrane (wetted briefly with methanol) in methanol-free transfer buffer (25 mM Tris, 192 mM glycine) at 100 mA constant current, at 4°C, with stirring. After transfer, the membrane was blocked at least one hour in 5% milk in Tris-buffered saline (TBS), then placed in a 4×4 inch resealable plastic zip lock bag, to which 2.5 ml 5% milk / TBS plus primary antibody was added in an even stream across the surface of the membrane. Excess air was released before sealing the bag, and then the antibody solution was massaged back and forth over the membrane for 1 minute, and incubated for 1 hour at RT. The membrane was rinsed briefly in 5% milk / TBS before a similar incubation step in HRP-conjugated secondary antibody. Two hours of vigorous washes in TBS on a shaker followed. The membrane was incubated with Western Lightning Plus-ECL detection reagents (Perkin Elmer #NEL104001EA), sandwiched between layers of Saran Wrap, and exposed to HyBlot CL film (Thomas Scientific, #1141J52). Primary antibodies used were as follows: mouse anti-GFP (Millipore Sigma), 1:2000; rabbit anti-HA (Cell Signaling), 1:2000; mouse anti-V5 (Thermo-Fisher), 1:1000; guinea pig anti-Syp (Davis lab, McDermott et al., 2014), 1:20,000; mouse anti-Tubulin (Cell Signaling #3873S), 1:500.

### Immunostaining

For anti-CycB staining, testes were dissected in 1xPBS, fixed in 1 mL ice-cold methanol (5 minutes), washed in 1 mL ice-cold acetone (2 minutes), and then washed in 1 mL room-temperature PBS + 0.1% Triton (PBSTr). Testes were blocked 30 minutes in 1 mL 3% BSA / PBSTr and incubated in anti-CycB (mouse F2F4, Developmental Studies Hybridoma Bank) overnight at 1:50 in 3% BSA / PBSTr at 4°C. After a one-hour wash in 1 mL 3% BSA / PBSTr, testes were incubated in secondary antibody (Alexa Fluor-conjugated donkey anti-mouse 488, 1:200, Molecular Probes) for 2 hours in the dark at room temperature. After two 10-minute washes in PBSTr, testes were transferred to a slide, where the liquid was removed and replaced with one drop of DAPI-containing Vectashield (Vector Labs). A 22×40 mm coverslip was placed on top and secured with nail polish. Samples were imaged within a few hours of mounting.

For all other antibodies, testes were dissected in 1xPBS in a 1.7-ml Eppendorf tube, fixed in 1 mL 4% formaldehyde/PBS for 30 minutes, washed briefly in 1 mL PBSTr, and permeabilized in 1 mL 0.3% sodium deoxycholate / 0.3% Triton-X / PBS for 30 minutes. After a quick wash in PBSTr, samples were processed through block, primary antibody, wash, secondary antibody, washes, and mounting, as described above. Primary antibodies: rabbit anti-Syp 1:200 (Rossi & Desplan 2020), anti-Tj 1:100 (guinea pig; a gift from D. Godt, University of Toronto, Canada), rabbit anti-HA (1:100) (Cell Signaling), and chicken anti-GFP 1:10,000 (Abcam #13970). Alexa Fluor-conjugated secondary antibodies: donkey anti-rabbit 488, anti-guinea pig 549, anti-chicken 488, and anti-rabbit 568 from Molecular Probes were used at 1:200.

### Single-molecule FISH

Testes were dissected from 1-2 day old flies in 1X PBS, fixed in 5% formaldehyde for 30 minutes, then permeabilized by incubation in PBS with 0.3% Triton X-100 and 0.6% sodium deoxycholate for 30 minutes at room temperature. After permeabilization we followed the conventional smFISH protocol in *Drosophila* (Trcek et al., 2017) while using the primary/secondary probe strategy from the inexpensive version (Tsanov et al., 2016; Gallicchio et al., 2021). Samples were mounted using 10 μl of ProLong™ Diamond Antifade Mountant with DAPI (Thermo-Fisher). Primary probes against *cycB* RNA (listed in Table S1) were designed using the Biosearch Technologies Stellaris probe Designer (version 4.2). To the 5ʹ end of each probe was added the Flap sequence TTACACTCGGACCTCGTCGACATGCATT. A secondary probe complementary to the Flap sequence was tagged with fluorophore CAL Fluor Red 610.

Imaging of smFISH samples was performed using a Leica SP8 Inverted Tandem Head confocal microscope and LAS X v3.5.7.23225 software (Cell Science Imaging Facility, Stanford University). Images were all taken at 40x magnification using the built-in tiling function to reconstruct the entire testis. The quantification of smFISH signal in Figures 5 and 9 was performed on three testes per genotype, then averaged to give the values shown. Relative fluorescence intensity was calculated for each testis by using FIJI to measure mean gray value (MGV) in squares of mid-stage and late-stage spermatocyte cytoplasm (5+ per testis) as well as squares of background staining (3+ per testis) from early spermatocytes (which have not yet turned on *cycB* from the spermatocyte promoter), then subtracting the average MGV of background from the average MGV of mid/late spermatocytes.

### Microscopy

Images from immunostaining and phase-Hoechst staining were captured by a Photometrics CoolSNAP CCD camera connected to a Zeiss Axioskop microscope, with fluorescence illumination provided by an X-Cite 120 excitation light source. Phase imaging was conducted on the above setup (Fig. 4A-D) or with a Spot RT3 CCD camera affixed to a Zeiss Axioskop microscope (Fig. 4G-I, Fig. 5D-F). For Fig. 5D-F, overlapping photos were taken at 20x and stitched together in Photoshop Elements. For scoring meiotic entry and exit (Fig. 4), nuclei were not considered round enough to count as “meiotic entry” if they were football- or teardrop-shaped, or if they had a lot of wobble when the focus shifted along the z axis. Cells were not scored as round spermatids if the mitochondrial derivative was smudgy / lacked a smooth circumference.

## ACKNOWLEDGEMENTS

We thank the Fuller lab and members of the Department of Developmental Biology for helpful discussions; the Davis lab (notably Ilan Davis, Lu Yang, and Darragh Ennis) for Syp-related conversation and their guinea pig anti-Syp antibody; and Claud Desplan for sharing rabbit anti-Syp antibody. The Stanford Cell Science Imaging Facility (CSIF) and Protein and Nucleic Acid Facility (PAN) were also instrumental to this work. We also thank the Bloomington Stock Center, Vienna Drosophila Resource Center, and the Drosophila Genomics Resource Center for making fly strains readily available. Finally, we would like to acknowledge FlyBase as an absolutely indispensable resource for this project and the *Drosophila* community in general.

## AUTHOR CONTRIBUTIONS

L.G. performed the smFISH experiments, including imaging. L.P. discovered the *syp* phenotype in an RNAi screen and determined which promoters/isoforms were enriched in testis and needed to be targeted by CRISPR. E.T. performed some of the early work on the Syp protein and its interactions. C.T. contributed to the molecular cloning for the Syp transgenes. C.C.B. did the remainder of the experimental work. C.C.B. and M.T.F. designed the study and wrote the paper.

---

**Figure S1.**
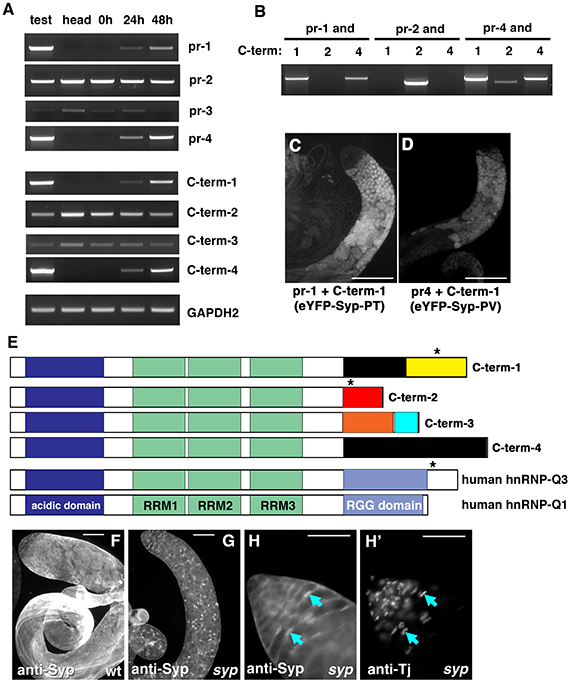
(A) RT-PCR of the four *syp* promoters (rows 1-4), the *syp* C-terminal variants (rows 5-8), and *GAPDH2* (last row) from cDNA from testis, head, and time-course testes (0h [*bam*], 24h, and 48h PHS). (B) RT-PCR from testis cDNAs for several pairwise combinations of promoters and C-termini, as indicated. (C,D) live fluorescence imaging of (C) eYFP-Syp-PT (promoter 1 + C-term-1) and (D) eYFP-Syp-PV (promoter 4 + C-term-1). Both reporter proteins localized to the nucleus in early spermatocytes (consistent with the NLS encoded in C-term-1) but largely shuttled to the cytoplasm in later spermatocytes. Scale bar: 200 μm. (E) Diagram of predicted Syp proteins, with two isoforms of human HNRNPQ at bottom. Black asterisks indicate nuclear localization signals (NLSes). The acidic domain is enriched for aspartic acid and glutamic acid residues; the RRM domains are classic RNA-binding domains; and the RGG domain in human HNRNPQ is enriched in arginine-glycine-glycine repeats. (F,G) anti-Syp (rabbit) on (F) wild-type and (G) *syp* testes. Scale bar: 100 μm. (H,H’) High-magnification view of the testis apical tip from *syp* male stained with (H) rabbit anti-Syp and (H’) anti-Tj. Arrows: somatic nuclei expressing both proteins. Scale bar: 50 μm (H,H’).

**Figure S2.**
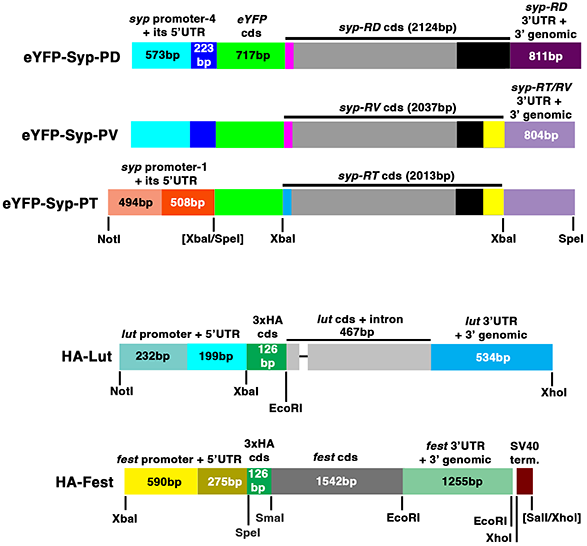
Diagram of new transgenes described in Materials and Methods. Not shown: V5-Lut, which is similar to HA-Lut but with a V5 coding sequence (42bp) in place of 3xHA.

**Table S1.**
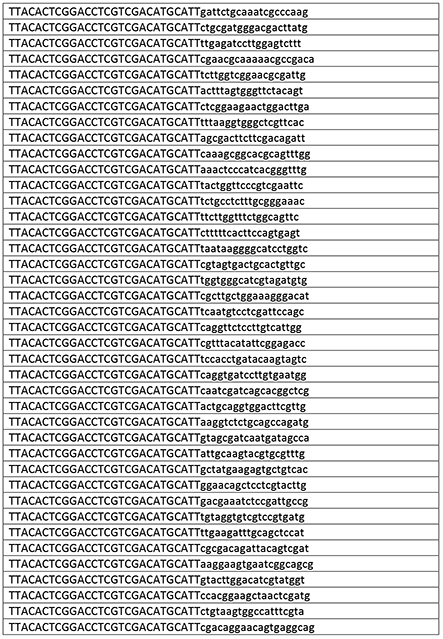
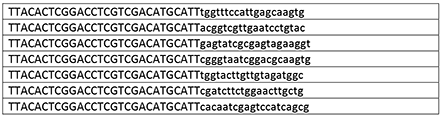
*cycB* primers for smFISH.

